# Post-capillary venules is the locus for transcytosis of therapeutic nanoparticles to the brain

**DOI:** 10.1101/2020.06.05.133819

**Authors:** Krzysztof Kucharz, Kasper Kristensen, Kasper Bendix Johnsen, Mette Aagaard Lund, Micael Lønstrup, Torben Moos, Thomas Lars Andresen, Martin Johannes Lauritzen

## Abstract

Treatments of neurodegenerative diseases require biologic drugs to be actively transported across the blood-brain barrier (BBB). To answer outstanding questions regarding transport mechanisms, we determined how and where transcytosis occurs at the BBB. Using two-photon microscopy, we characterized the transport of therapeutic nanoparticles at all steps of delivery to the brain and at the nanoscale resolution *in vivo*. Transferrin receptor-targeted nanoparticles were taken up by endothelium at capillaries and venules, but not at arterioles. The nanoparticles moved unobstructed within endothelial cells, but transcytosis across the BBB occurred only at post-capillary venules, where endothelial and glial basement membranes form a perivascular space that can accommodate biologics. In comparison, transcytosis was absent in capillaries with closely apposed basement membranes. Thus, post-capillary venules, not capillaries, provide an entry point for transport of large molecules across the BBB, and targeting therapeutic agents to this locus may be an effective way for treating brain disorders.

**HIGHLIGHTS:**

- Integration of drug carrier nanotechnology with two-photon microscopy *in vivo*
- Real-time nanoscale-resolution imaging of nanoparticle transcytosis to the brain
- Distinct trafficking pattern in the endothelium of cerebral venules and capillaries
- Venules, not capillaries, is the locus for brain uptake of therapeutic nanoparticles

## INTRODUCTION

The blood-brain barrier (BBB) is impermeable to most blood-borne substances (Abbott et al., 2010). The paracellular route across the BBB is limited by junctional complexes between brain endothelial cells (BEC). Diffusion across BEC is possible, but restricted to low-molecular weight hydrophobic compounds, and most lipophilic drugs show negligible brain uptake because of the rapid outward transport of xenobiotics by BEC efflux pumps (Mollgard et al., 2017; Pardridge, 2012). Large molecules can enter the brain, but only via vesicular transport, i.e. transcytosis. However, this route is highly selective and strongly suppressed by recently identified homeostatic mechanisms (Ben-Zvi et al., 2014; De Bock et al., 2016; Janiurek et al., 2019; Tietz and Engelhardt, 2015). Consequently, while the low BBB permeability protects the brain against circulating pathogens, it also precludes more than 98% of neuroprotective compounds from reaching the brain, rendering central nervous system (CNS) disorders resistant to most conventional therapies (Banks, 2016; Pardridge, 2002, 2012).

Drug delivery systems that adapted the receptor-mediated transcytosis (RMT) to shuttle large therapeutic cargo across the BBB are currently at the forefront of the modern pharmacology (Lajoie and Shusta, 2015; Pardridge, 2020; Pulgar, 2018). Coupling carrier molecules to RMT ligands may enhance delivery of therapeutics to the brain, and the flagship RMT ferrying receptor in the brain is the transferrin receptor (TfR) (Johnsen et al., 2019b; Pardridge, 2002, 2020). Nanoparticles, such as liposomes, are versatile drug carriers that can encapsulate a variety of xenobiotics, and functionalized with TfR ligands represent a promising drug delivery approach with diverse therapeutic opportunities (Johnsen et al., 2019b; Pardridge, 2020). The translational potential of TfR-targeted nanoparticles has been tested in preclinical models of Parkinson’s, Alzheimer’s, Huntington’s disease, brain cancer, and stroke, but the level of drug delivery levels rarely meet the thresholds for clinical significance due to failure in nanoparticle transcytosis (Johnsen et al., 2019b).

To address this issue, we examined how cerebral vessels handle clinically relevant nanoparticles in the living, intact brain. We employed two-photon microscopy in anesthetized and awake mice to characterize the pharmacokinetics of TfR-targeted nanoparticles at the BBB, intracellular trafficking patterns in BECs, and transcytosis into the brain parenchyma using real-time *in vivo* imaging at nanoscale resolution. We report that TfR-targeted nanoparticles bind efficiently to BECs in venules and capillaries, but are absent in arterioles. Despite the highest uptake by BECs in capillaries, we found that transcytosis occurs only in post-capillary venules that, in contrast to capillaries, are surrounded by a large perivascular space that can accommodate nanoparticle-sized elements (Engelhardt et al., 2017). Our observations indicate that post-capillary venules mediate brain uptake of large drug carriers, which should be considered in the design of next-generation nanoparticles for the treatment of brain disorders.

## RESULTS

### Two-photon imaging of BBB-targeted nanoparticles *in vivo*

To investigate how drug nanocarriers interact with the BBB *in vivo*, we used two-photon fluorescence microscopy in mice. Following microsurgery, the brain was imaged via a cranial window over the somatosensory cortex (Figures 1A and 1B). The nanoparticles were designed to resemble clinically approved nanoliposome formulations (Barenholz, 2012), consisting of a distearoylphosphatidylcholine(DSPC)/cholesterol lipid bilayer surrounding an aqueous lumen and with a PEGylated surface to ensure stability in the blood (Figure 1C, Table S1). To enable targeting to the brain via TfR, we coupled the nanoparticles to immunoglobulin G (IgG) monoclonal anti-TfR antibody clone RI7217 (Lee et al., 2000). Prior to two-photon imaging experiments, we validated the nanoparticle TfR targeting system by encapsulating BBB-impermeable cisplatin into nanoparticles (Table S1), and injecting the nanoparticles into the bloodstream. The traces of cisplatin were detected in the brain only for nanoparticles functionalized with RI7217, but not stealth (no targeting antibody), or isotype IgG (no TfR recognition) antibodies (Figure 1D).

**Figure 1.**
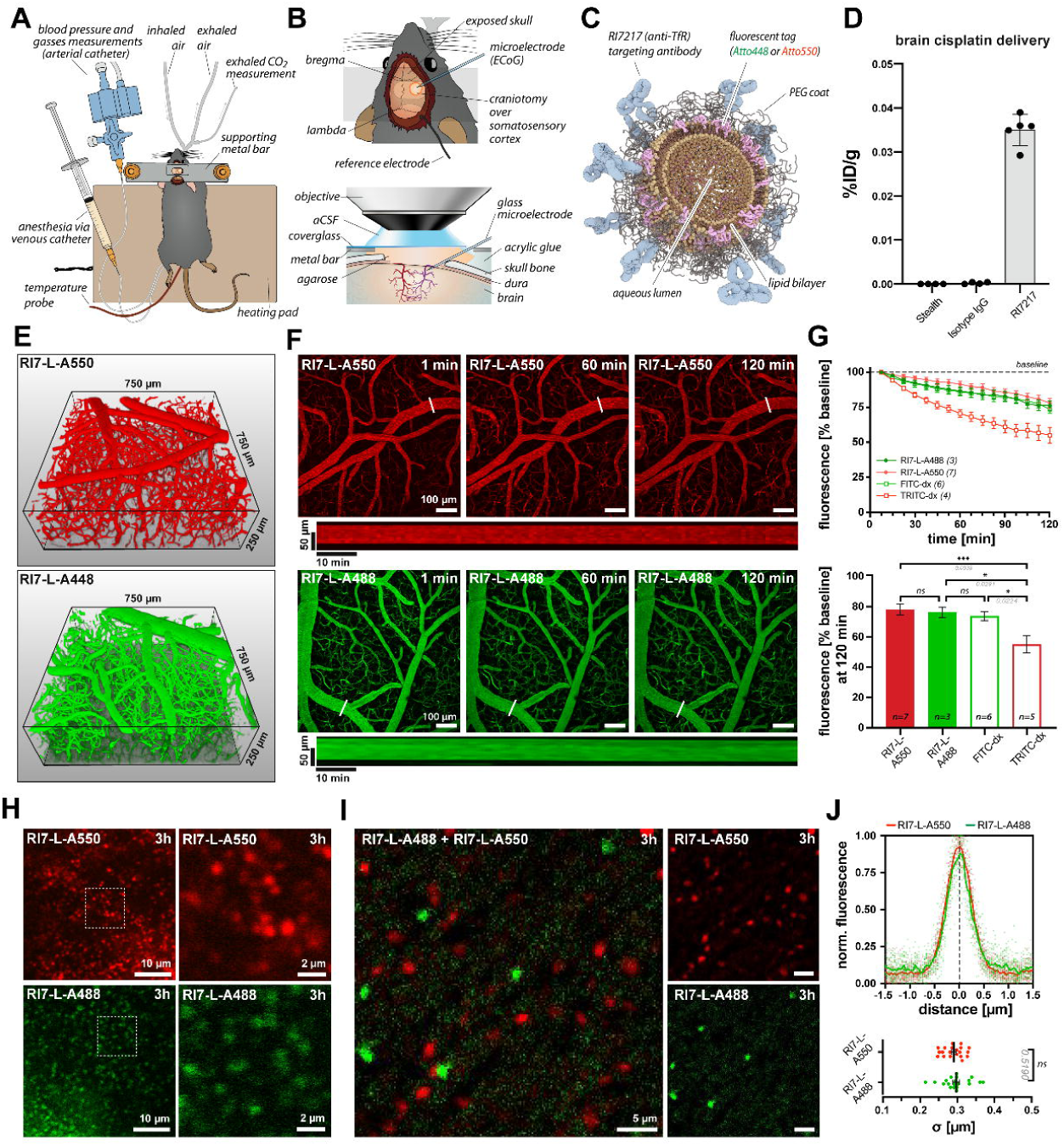
Two-photon imaging of liposome nanoparticles *in vivo*. **A)** Mouse after the preparative surgery for acute imaging. **B)** Features of an acute cranial window. **C)** Principal components of targeted nanoparticles used in the study. **D)** RI7217-functionalized nanoparticles drive accumulation of cisplatin payload in the brain, in contrast to nanoparticles functionalized with isotype IgGs or devoid of antibody functionalization (stealth liposomes). %ID/g is percentage of injected dose of encapsulated cisplatin per gram tissue; *n*=4 mice. **E)** 3D reconstruction of the brain microvasculature delineated by circulating nanoparticles. **F)** Time-lapse images of blood-circulating nanoparticles following injection into the bloodstream. Long panels are kymographs of fluorescence at demarked lines. See also Movie S1. **G)** Upper panel: nanoparticle fluorescence signal over time. Lower panel: fluorescence signal from RI7-L-A550 and RI7-L-A488 nanoparticles after 2 h of continuous imaging matches the stability of FITC-dx and outperforms TRITC-dx, commonly used in imaging studies (ANOVA). *(n)*=number of mice **H)** Laser-extravasated nanoparticles into brain parenchyma after 3 h after injection into the circulation. Nanoparticles retained their discrete and homogeneous appearance. **I)** Laser-extravasated RI7-L-A488 and RI7-LA550 into the brain parenchyma 3 h after injection into the circulatiion. No liposome fusion or aggregation occured when present in circulation or in the brain. **J)** Average of fluorescence profile plots of nanoparticles laser-extravasated to the brain 3 h after injection into the circulation. The average standard deviation (σ) of signal distribution did not differ between distinctively labeled nanoparticles. *(n)*=number of nanoparticles. *p<0.05; ***p<0.001, *ns=*non-significant. Data are means±SEM.

To enable detection in subsequent two-photon imaging experiments, we fluorescently labeled the nanoparticle lipid bilayer with either Atto488 or Atto550 (Figure 1C). Both the Atto550- and Atto488-labeled nanoparticles (RI7-L-A550 and RI7-L-A488) were administered by single bolus injection into the bloodstream (70 nmol_lipid_/g_animal_) and were subsequently imaged in the brain volume containing pial arterioles and venules, penetrating arterioles, ascending venules, pre- and post-capillary vessels, and capillaries (Figure 1E). The blood-circulating nanoparticles exhibited high fluorescence stability over time, indicating a low filtration rate by peripheral organs (Figures 1F and 1G, Movie S1). Nanoparticles retained their single-particle appearance without agglomeration or fusion, as ascertained by laser-extravasation of nanoparticles from the bloodstream into the brain parenchyma after 3 h in circulation (Figures 1H and 1I). A small fraction of nanoparticles was sequestered by leukocytes, but without adverse effects on the brain and systemic physiology that could compromise the BBB (Figure S1, Movie S2).

Nanoparticles in the brain *in vivo* exhibited properties of a point source signal with a Gaussian distribution peak denoting the center location of the nanoparticle. The fluorescence distribution standard deviation (σ) was 0.290±0.006 µm and 0.296±0.008 µm for RI7-L-A550 and RI7-L-A488, respectively, and did not differ between the two distinctively labeled nanoparticle types (n=20, p=0.5910, t-test) (Figure 1J). In subsequent experiments, the nanoparticles were considered spatially separated when their peak of fluorescence exceeded 2σ from other sources of fluorescence, i.e., vessel lumen, endothelium, or other nanoparticles.

Thus, the nanoparticles fulfilled all necessary requirements for *in vivo* imaging: efficient two-photon excitation and emission of fluorescence with the ability to resolve single nanoparticles, high structural stability in the circulation, and no detrimental effects on the brain and systemic physiology.

### Targeted nanoparticles selectively associate to capillaries and venules

To delineate vessel lumen, we used FITC-dextran (FITC-dx) injected subsequently to the nanoparticles. The blood-circulating nanoparticles were rapidly recruited to the endothelium, and at 2 h post-injection, appeared as numerous punctae at vessel walls of the capillaries, and of the pial (Figure 2A), ascending, and post-capillary venules (Figures 2B–2E), but were absent from the arterial branches of the brain microvasculature (Figure 2F). The association was driven by interactions of the TfR-Ab with the TfR at the BEC surface and not by stalling of nanoparticles in the vessel lumen, as neither antibody-lacking stealth nanoparticles (Sth-L-A550)(Figures S2A and S2B) nor isotype IgG-functionalized nanoparticles that lack TfR recognition (IgG-L-A550) associated to BECs (Figures S2C and S2D). The nanoparticle association was independent of the type of fluorescent tag, as RI7-L-A488 exhibited the same targeting properties as RI7-L-A550 (Figures S3A–E).

**Figure 2.**
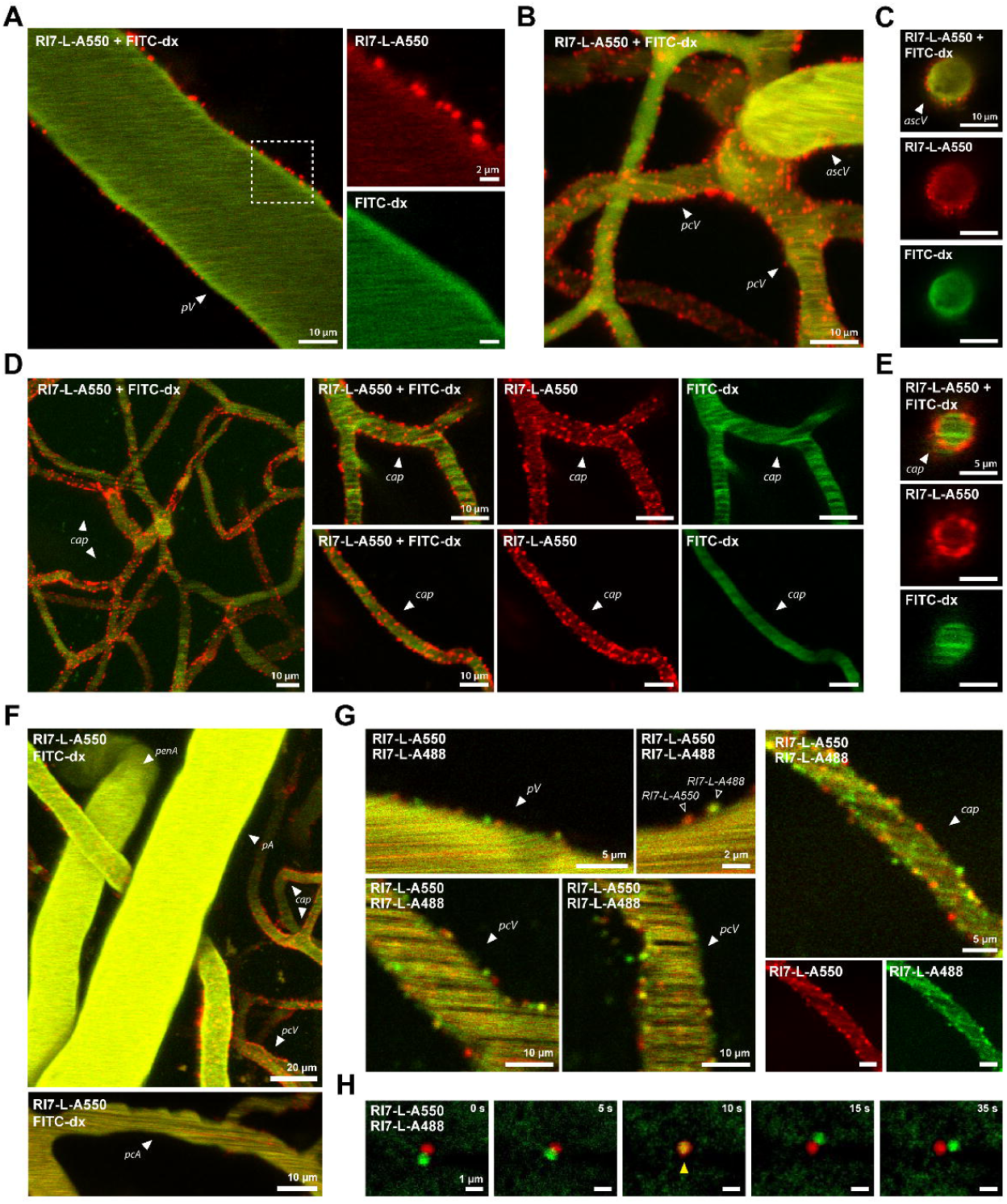
Robust association of targeted nanoparticles to capillaries and post-capillary venules. **A–E)** *In vivo* images of RI7-L-A550 nanoparticles (red) *in vivo* 3 h after injection to the circulation. Co-injected circulating FITC-dx delineates vessel lumen (green). Liposomes associate with vessel walls at pial, ascending venules, post-capillary venules, and capillaries. **F)** No association of liposomes to an arterial branch of the brain microvasculature. **G)** Co-injection of both RI7-L-A550 and RI7-L-A488 revealed discrete punctae of both nanoparticles at the vessel walls. **H)** Time-lapse images of laser-extravasated nanoparticles in brain parenchyma. The proximity of nanoparticles causes overlapping fluorescence signal. *pV=*pial venule; *ascV=*ascending venule; *pcV=*post-capillary venule; *cap=*capillaries; *penA*=penetrating arteriole; *pA=*pial arteriole.

Lastly, mice co-injected with both nanoparticles exhibited distinct labeling at the vessel walls from both RI7-L-A488 and RI7-L-A550, attesting that the observed punctae represented single nanoparticles (Figure 2G). Scarce presence of merged signals was attributed to overlapping fluorescence of nanoparticles separated by a distance smaller than the diffraction limit of the microscope (Figure 2H).

### Vessel type determines association density and kinetics

We isolated BEC-associated from blood-circulating nanoparticles by excluding the signal of circulating nanoparticles that colocalized with FITC-dx in the vessel lumen (Figure 3A). Three-dimensional (3D) reconstruction revealed a spatially heterogeneous association of nanoparticles across the vascular tree (Figure 3B). At 2 h post-injection, the highest density, i.e., number of nanoparticles at the vessel surface, was at BECs of capillaries and venules with an exponential decline in ascending and pial venules (Figure 3D). This observation was consistent with the kinetics of association being fastest in capillaries and slowest in pial venules (Figures 3E, S3F, and S3G, Movie S3). Of note, nanoparticle binding was ongoing even after 2 h post-injection, indicating that the cellular pool of available TfRs was not saturated at this time (Figure 3E).

**Figure 3.**
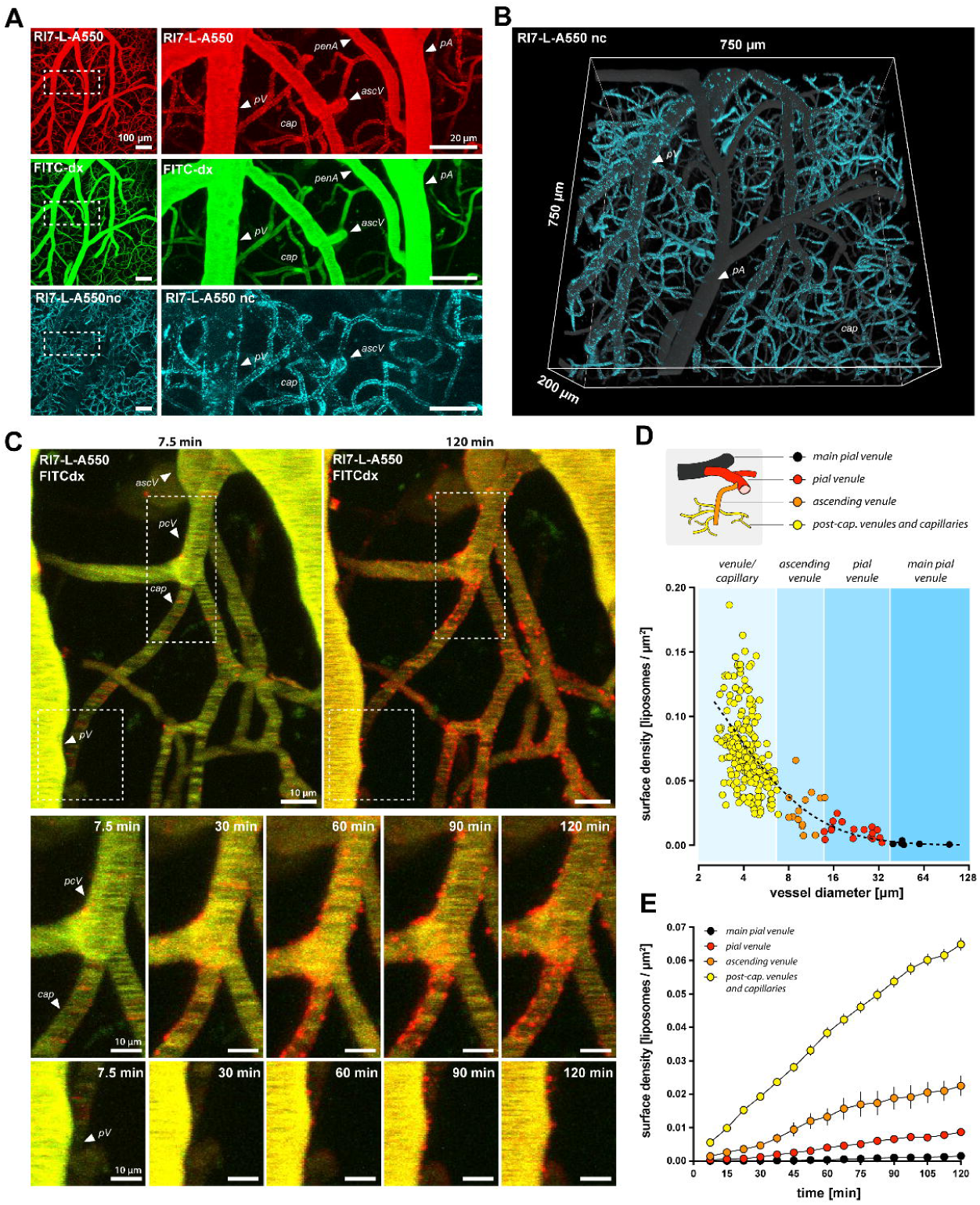
Nanoparticle association density and kinetics are determined by vessel type. **A)** The difference between RI7-L-A550 (red) and FITC-dx (green, vessel lumen) fluorescence signal used to show the fraction of non-circulating nanoparticles that contour vessel boundaries (RI7-L-A550nc, cyan). **B)** 3D reconstruction of the nanoparticles associatiod to endothelium 2 h post-injection. The recuruited (non-circulating) nanoparticles (RI7-L-A550nc, cyan) are superimposed on the signal from liposomes circulating in the bloodstream (gray). **C)** Upper panel: brain microvasculature following co-injection of RI7-L-A550 nanoparticles (red) and FITC-dx (vessel lumen, green) into circulation. Squares indicate areas magnified in lower panels. Lower panels: association of nanoparticles over time. See also Movie S3. **D)** Nanoparticle distribution 2 h post-injection. Each data point is a single vessel. Blue areas indicate vessel segments. Dashed line demarks lognormal distribution trendline. Inset illustrates vessel classification. **E)** Nanoparticles associated most rapidly with capillaries and most slowly with large venules. Surface density=# of nanoparticles per µm^2^ vessel wall area. *pV=*pial venule; *ascV=*ascending venule; *pcV=*post-capillary venule; *cap=*capillaries; *pcA=pre-capillary arteriole; penA*=penetrating arteriole; *pA=*pial arteriole. Data are means±SEM.

### Most of associated nanoparticles are internalized by endothelium

To determine the type of interaction between nanoparticles and BEC, i.e. the ratio between internalized and adhering nanoparticles, we used transgenic mice expressing cytosolic green fluorescent protein (GFP) in the endothelium (Tie2-GFP; see Methods). This allowed for a comprehensive assessment of endothelium morphology with unprecedented details in the living brain (Figures 4A and S4). We could discern the distinct patterns of nuclei and endothelial cell contact sites (Figure 4B), and the fluorescence signal was compatible with imaging of the nanoparticles (Figure 4C).

**Figure 4.**
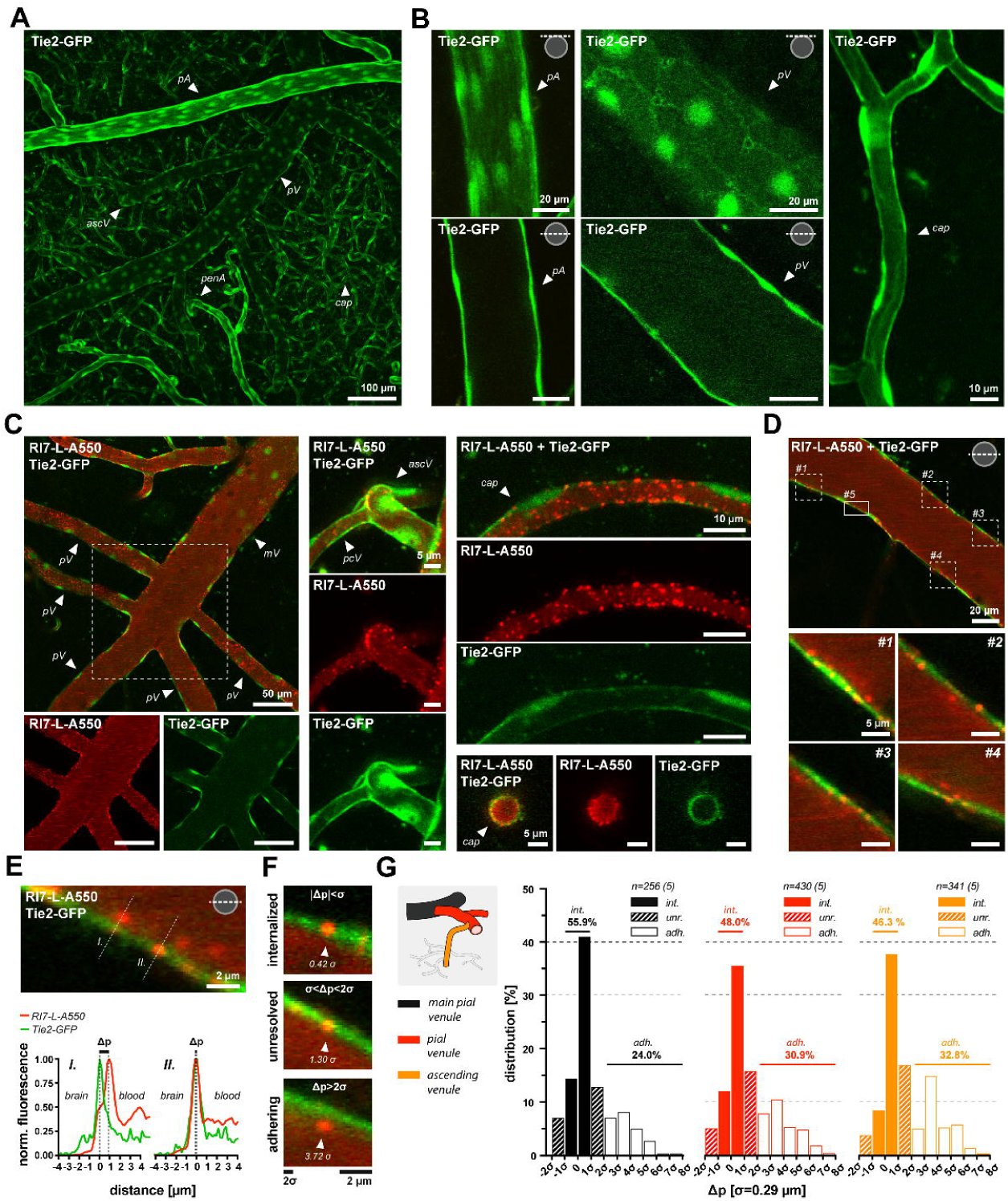
Prevalence of internalized nanoparticles in the brain vasculature. **A)** Brain endothelium *in vivo* imaged in Tie2-GFP mice. **B)** Detailed morphology of endothelial cells *in vivo* with noticeable cell boundaries and bright nuclei. **C)** Simultaneous imaging of the brain endothelium and nanoparticles 2 h after RI7-L-A550 injection into circulation. **D)** Pial venule with a clear presence of nanoparticles. Boxes 1–4 represent magnified areas on the right panels. #5 is the magnified area presented in E). **E)** Upper panel: nanoparticles (red) at two distinct locations (*I*., *II*.) in relation to the endothelium (Tie2-GFP, green). Dashed lines indicate the axes of fluorescence profiles in the lower panel. Lower panel: fluorescence profiles of a nanoparticles with low (*I*.) and high signal overlap (*II*). Δp demarks signal peaks separation. **F)** Examples of internalized (|Δp|<σ), unresolved (σ<|Δp|<2σ), and adhering (|Δp|>2σ) nanoparticles. σ=0.29 µm. Scale bars are provided in both σ and µm. Numbers at arrowheads show Δp value in σ units. **G)** Histogram of percentage distribution of internalized (*int*.), unresolved (*unr*.), and adhering (*adh*.) liposomes. Inset illustrates vessel type division. Values over horizontal lines are the sum of underneath bins. *n*=number of liposomes, values in brackets are the number of mice. Histogram bin=σ=0.29 µm. Image insets denote the position of imaging plane relative to a vessel perimeter. *mV=main pial venule; pV=*pial venule; *ascV=*ascending venule; *pcV=post-capillary venule; cap=*capillaries; *penA*=penetrating arteriole; *pA=*pial arteriole.

At 2 h post-injection, TfR-coated nanoparticles exhibited a non-uniform distribution in relation to the vessel wall, appearing either to be internalized or to adhere to the luminal surface of the endothelium (Figure 4D). For each analyzed vessel, we aligned the imaging plane with the center of the vessel radial symmetry (the broadest vessel cross-section), and for each nanoparticle, we extracted the fluorescence profile plot with corresponding endothelium signal (Figure 4E). As a measure of separation, we used the distance between peaks (Δp). The precise spatial localization of nanoparticles is non-trivial because of the diffraction limit, therefore we defined the boundary conditions characterizing nanoparticle location using the previously calculated standard deviation (σ) of Gaussian fluorescence distribution (σ=0.29 µm; Figure 1J). Using conservative inclusion criteria, the nanoparticles were categorized as *internalized* when separated from endothelium peak signal only by |Δp|<σ (high fluorescence overlap) and as *adhering* when separated from the endothelium peak signal by at least 2σ, i.e., |Δp|>2σ (minimal fluorescence overlap). The third group consisted of nanoparticles that could not be categorically classified into adhering or internalized and represented the intermediate group (i.e., *unresolved*, σ ≤ |Δp| ≤ 2σ). Our boundary conditions were further attested by visual inspection of nanoparticles (Figure 4F). Given a wide signal point spread function along the Z-axis (depth), we did not analyze capillaries because of their small diameter and high vessel curvature. At 2 h post-injection, the internalized fractions were 55.9%, 48.0%, and 46.3% of total nanoparticles at the endothelium for main pial venules, pial venules, and ascending venules, respectively (Figure 4G). The corresponding adhering fraction of nanoparticles was 24.0%, 30.9%, and 32.8%, while the unresolved fraction was 20.1%, 21.1%, and 20.8%. This indicates that, on average, at least half of the nanoparticles recruited to vessel walls were internalized by endothelium. Noteworthy, within the *unresolved* fraction, we detected nanoparticles with Δp < -σ, with prevalences of 7.1%, 5.2%, and 3.8% for main pial venules, pial venules, and ascending venules, respectively. These nanoparticles were located in proximity to the basolateral membrane of BEC, and their prevalence increased with venule diameter.

### Directional movement of nanoparticles inside endothelium

Once internalized, the nanoparticles exhibited a relatively high degree of movement over short distances (Figure 5A, Movie S4). The nanoparticles were tracked at 2-3 h post-injection with respect to large vessels, i.e. pial venules, and small vessels, i.e. post-capillary venules with capillaries (Movie S5). We omitted ascending venules because of their varying angled orientation to the imaging plane. From each analyzed vessel, we selected 10 nanoparticles that remained in the imaging plane throughout 30 mins of continuous imaging (time interval between frames=30 s) (Figure 5B). The traces were aligned to the point of origin (Figure 5C), and to avoid underestimation of movement in planar *(x,y)* coordinates because of vessel curvature, we projected each trace to a vector *(v)* aligned with a vessel symmetry axis and blood flow direction, both independent from circumference curvature (Figure 5D).

**Figure 5.**
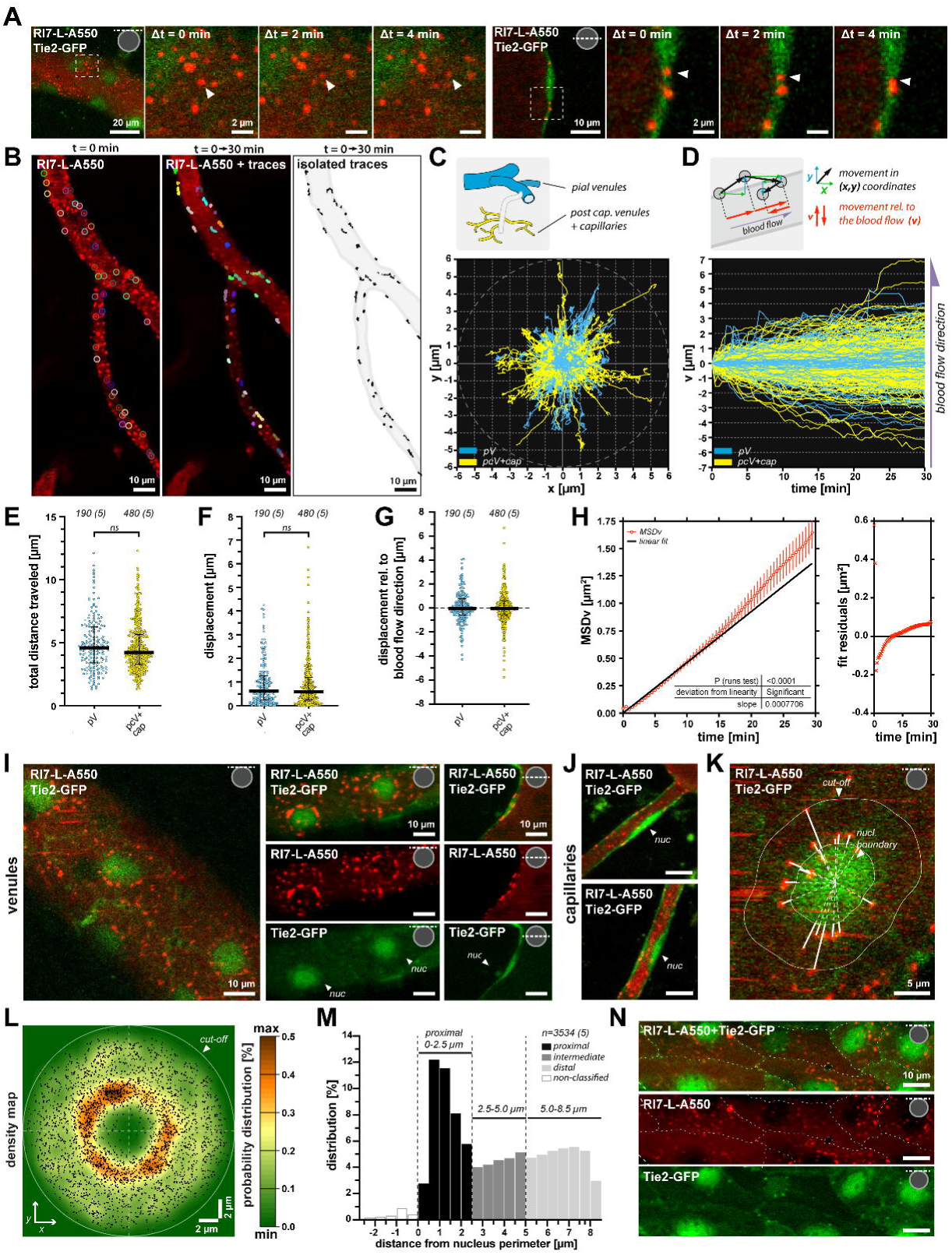
Nanoparticles exhibit directional movement in all vessel types, but distinct subcellular distribution. **A)** Time-lapse images of nanoparticle movement in large venules. *Left panel:* vessel surface scan. *Right panel:* vessel cross-section scan. Arrowheads indicate the movement of a nanoparticle. See also Movie S4 **B)** Tracking of nanoparticles. *Left panel:* circles outline traced nanoparticles. *Middle panel:* movement traces during 30 min of continuous time-lapse imaging. *Right panel:* isolated movement traces (black) with contours delineating the microvessel (gray). See also Movie S5 **C)** Movement traces in *x,y* (Cartesian) coordinate system with respect to a vessel type. (0,0) corresponds to the initial position of a nanoparticle. **D)** Translation of the movement in *x,y* coordinates into the movement relative to the blood flow direction *v. v>*0 indicates the movement along the blood flow, and *v*<0 is against the blood flow direction. **E-F)** Vessel type does not influence the total distance traveled and the displacement (end-position) of a nanoparticle. Time span=30 min. *n*=number of nanoparticles, values in brackets are the number of mice. **G)** Blood flow direction does not affect the direction of movement of nanoparticles in large (pial venules) and small microvessels (post-capillary venules and capillaries). **H)** MSD analysis showing significant deviation of MSD(t) from linearity (Wald– Wolfowitz runs test). Solid line shows least squares linear regression fit. **I)** Nanoparticles preferentially distributed to perinuclear areas of venules. **J)** No preferential distribution in capillaries. **K)** Calculation of nanoparticles location (bold solid lines) in relation to nucleus perimeter (solid edges). Distance lines meet at the geometric center of a nucleus (dashed lines). **L)** Density map of nanoparticle distribution in relation to the geometric center of the nucleus. Kernel=2σ. The heat-map represents the probability of liposome presence at a given coordinate. **M)** Nanoparticle distribution in relation to nucleus boundary (bin=0.5 µm). Numbers over horizontal lines are the sum of underneath bins. Empty bars are liposomes overlapping with nucleus signal *(non-classified). n*=number of liposomes; number in brackets indicates number of animals. **N)** Nanoparticles do not distribute to the endothelial cells perimeter and contact sites. Dashed lines indicate cell boundaries. Insets in images denote the position of imaging plane relative to a vessel perimeter. *pV=*pial venule; *pcV=*post-capillary venule; *cap=*capillary; *nuc*=nucleus; *MSDv=*mean squared displacement in *v* coordinate. E–G) Data are medians with interquartile ranges (IQR). H) Data are means±SEM.

The total distance traveled along the vessel axis did not differ between nanoparticles in large venules, and post-capillary venules with capillaries (p=0.1672, Mann– Whitney; Figure 5E). Similarly, the end-position of the nanoparticle relative to the origin was the same for all vessel types (p=0.6694, Mann–Whitney)(Figure 5F). The movement did not occur preferentially along or against the blood flow, as in all vessel types, the median displacement relative to the blood flow direction approximated 0 (Figure 5G). Of note, nanoparticles exhibited motility even in capillaries with stalled blood flow (Movie S6). Thus, neither cell morphology, nor differences in blood flow velocities (Santisakultarm et al., 2012), nor blood flow direction influenced the intracellular movement of the nanoparticles.

To determine whether intracellular trafficking of nanoparticles occurs by random movement (e.g., diffusion) or is directional, we performed mean squared displacement (MSD) analysis (see Methods). MSD analysis was performed along the vessel axis *v* (Figure 5H) and MSD(t) significantly departed from linearity (p<0.001, n=673 nanoparticles, Wald–Wolfowitz runs test). This indicates that the movement in the endothelium contains a significant directional component. This suggested that the nanoparticles might be directed towards specific cell regions, which we assessed in the subsequent experiment.

### Nanoparticles distribute to endothelial cell perinuclear area in venules but not in capillaries

Upon internalization, the nanoparticles localized over time to the perinuclear region of endothelium cells in venules, but not in capillaries (Figures 5I and 5J). To quantify the spatial distribution of nanoparticles in venules, we measured the distances from the geometric center of the nucleus to the nanoparticles (Figure 5K). Our kernel density map shows that the highest probability for finding a nanoparticle in venules was in proximity to the nucleus (Figure 5L), at distances of 0.5–2.5 µm from the nucleus boundary and with numbers decreasing at intermediate (2.5–5 µm) and distal (>5 µm) regions (Figure 5M). Of note, neither vessel type exhibited clustering of nanoparticles at the endothelium perimeter (Figure 5N). This indicated that nanoparticles do not stall in the cytosol areas that contain dense cytoskeleton elements that support cell contact sites, and that nanoparticles do not wedge between adjacent endothelial cells.

### Nanoparticles are transcytosed to the brain, but only at venules

The possibility of large nanoparticle transcytosis across the BBB is disputed (Clark and Davis, 2015; Lindqvist et al., 2016; van Rooy et al., 2011; Wiley et al., 2013). Here, we detected nanoparticles present at the basolateral side of the endothelium in both capillaries and venules (Figure 6A). This could result from lateral displacement of nanoparticles, i.e. movement within endothelium along the vessel (Figure 6B). However, in contrast to capillaries, nanoparticles in venules exhibited translocation normal to the vessel lumen (Figure 6C). We observed that nanoparticles that associated with the vessel wall slowed down, but once transcytosed, they rapidly accelerated into the perivascular space (Figure 6D, Movie S7). This occurred only in pial and post-capillary venules and not in capillaries, indicating that the vessel type crucially determines successful transcytosis. Of note, we detected rare events in which blood-borne cells that sequestered nanoparticles crossed the BEC in venules (Movie S8), consistent with previous studies of immune-cell trafficking in the brain (Engelhardt et al., 2017).

**Figure 6.**
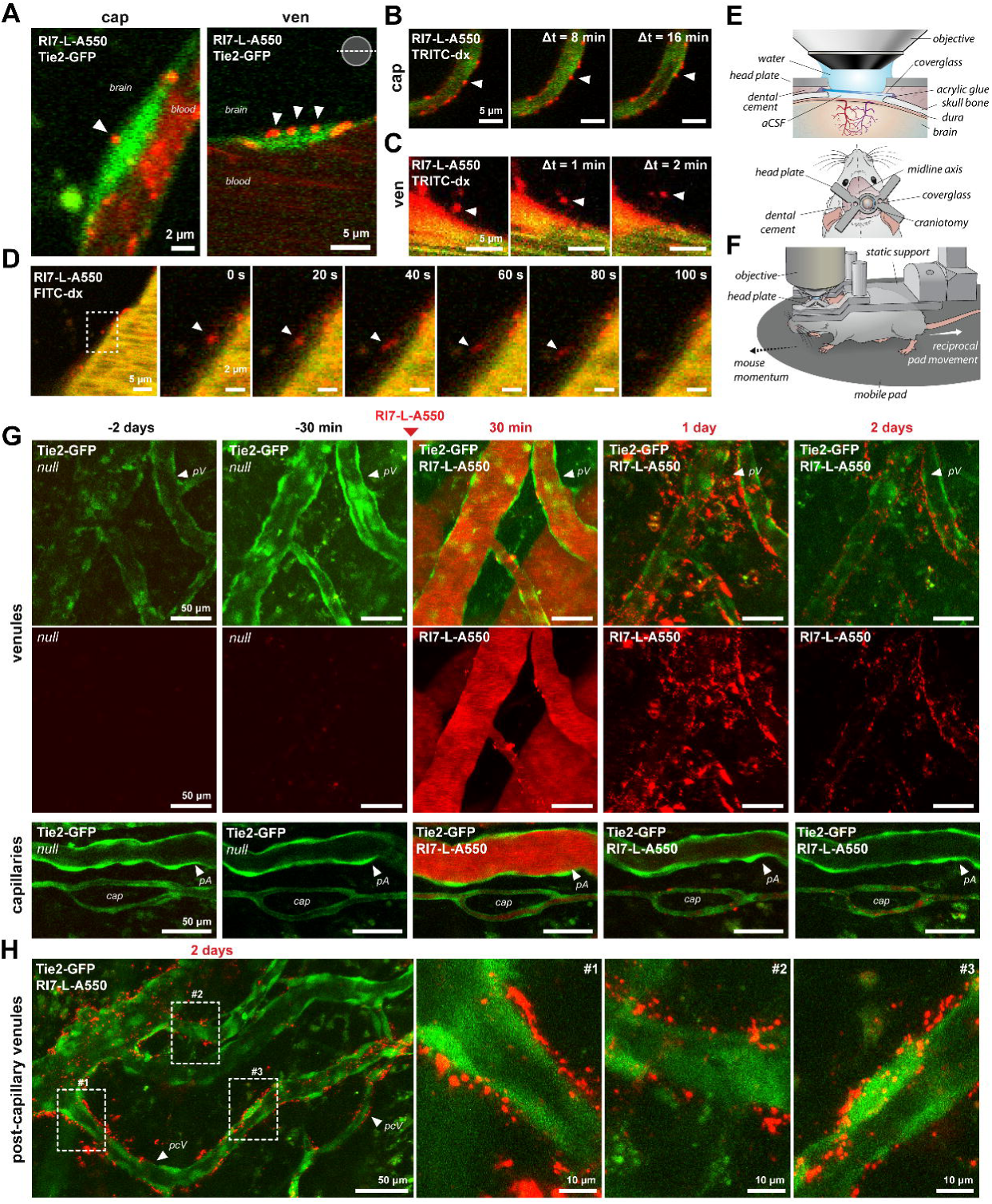
Post-capillary venules is the locus for TfR-mediated transcytosis. **A)** Internalized nanoparticles located at the basolateral side of endothelium in a capillary and a pial venule. **B)** Lateral nanoparticle movement in capillaries. **C)** Movement of a nanoparticle perpendicular to the vessel lumen. **D)** Nanoparticle crossing the endothelium into the perivascular space in venules. The images show a nanoparticle accelerating in the perivascular space. See also Movie S7. **E)** Schematic drawing of a mouse head and craniotomy for chronic two-photon imaging. **F)** Microscope stage with movement-unrestricted awake animal. The objective is stationary, and the air-pressurized pad reacts reciprocally to the mouse movement. **G)** Time course of nanoparticle penetration to brain parenchyma. Images are Z-stack maximum intensity projection. *Upper rows:* Nanoparticles enter the brain at post-capillary venular segments; *Lower row:* No nanoparticle transcytosis to the brain at capillaries. *null=*fluorescence signal prior to nanoparticles injection. **H)** High-resolution images of nanoparticles upon transcytosis in pial, ascending and precapillary venules. *pA=*pial artery; *pV=*pial venule; *pcV=*post-capillary venule; *cap=*capillary.

Continuous high-frequency time-lapse imaging required to capture transcytosis events may potentially damage the BBB, leading to extravasation of the blood-borne components to the brain (Choi et al., 2011). Given this possibility, along with the rarity of transcytosis events and limited span of acute imaging experiments (∼4 h), we next performed long-term two-photon imaging on awake and mobile mice with chronic cranial window implants (Figures 6E and 6F; see Methods). We imaged the brains of awake Tie2-GFP mice 10-day post-surgery, to identify the cortical areas that contained large venules (Figure 6G). We then reassessed the same loci 2 days later, and subsequently, injected RI7-L-A550 into the bloodstream (50 µL 70 nmol_lipid_/g_animal_). Next, we imaged the animals again (30 min post-injection) to ensure that the endothelium remained structurally intact and examined the same region at 1 and 2 days post-injection (Figure 6G). Already at 1 day post-injection, the animals exhibited no significant presence of nanoparticles in the circulation, indicating effective clearance of RI7-L-A550 from the bloodstream. However, at this timepoint we detected substantial amounts of nanoparticles on the abluminal side of the endothelium in the brain parenchyma surrounding venules. This fraction of nanoparticles significantly decreased between 1 and 2 days post-injection but was still abundant, indicating high retention (Figure 6G and H). In contrast to pial, ascending, and post-capillary venules, nanoparticles failed to transcytose to the brain tissue close to capillaries and the arterial branches of the microvasculature (Figure 6G). This confirmed that transcytosis of nanoparticles across the BBB occurs at venules and not at other segments of the brain microvasculature.

## DISCUSSION

Understanding the pharmacokinetics of drug-carrying nanoparticles at the BBB *in vivo* is crucial for future drug delivery strategies. For nearly two decades, drug delivery studies have faced methodological challenges to determine how nanoparticles may penetrate the BBB. Our current knowledge on nanoparticle-based drug delivery across the BBB is deduced from chemically processed tissue and biodistribution studies, or whole-brain imaging techniques (Cabezon et al., 2015; Johnsen et al., 2019b; Pardridge, 2012). Moreover, the microanatomy of brain tissue surrounding microvessels vastly differs among arterioles, capillaries, and venules (Engelhardt et al., 2017), but how this heterogeneity of CNS vascular anatomy defines distinct BBB compartments is unclear. Here, using two-photon microscopy, we provide direct insight into nanoparticle delivery through all stages of endothelial vesicular transport: binding, uptake, intracellular trafficking, and exocytosis in the living brain. We report that clinically relevant formulations of TfR-targeted nanoparticles rapidly associate with the BECs, with preferential distribution to capillaries. Upon internalization, nanoparticles exhibit movement kinetics independent of the vessel type, and with subcellular distribution profiles that differ between venules and capillaries. Despite the highest density of nanoparticles in capillaries, transcytosis across the BBB *in vivo* occurs only in venules (Figure 7), the same vascular segment that is involved in immune-cell trafficking (Engelhardt et al., 2017).

**Figure 7.**
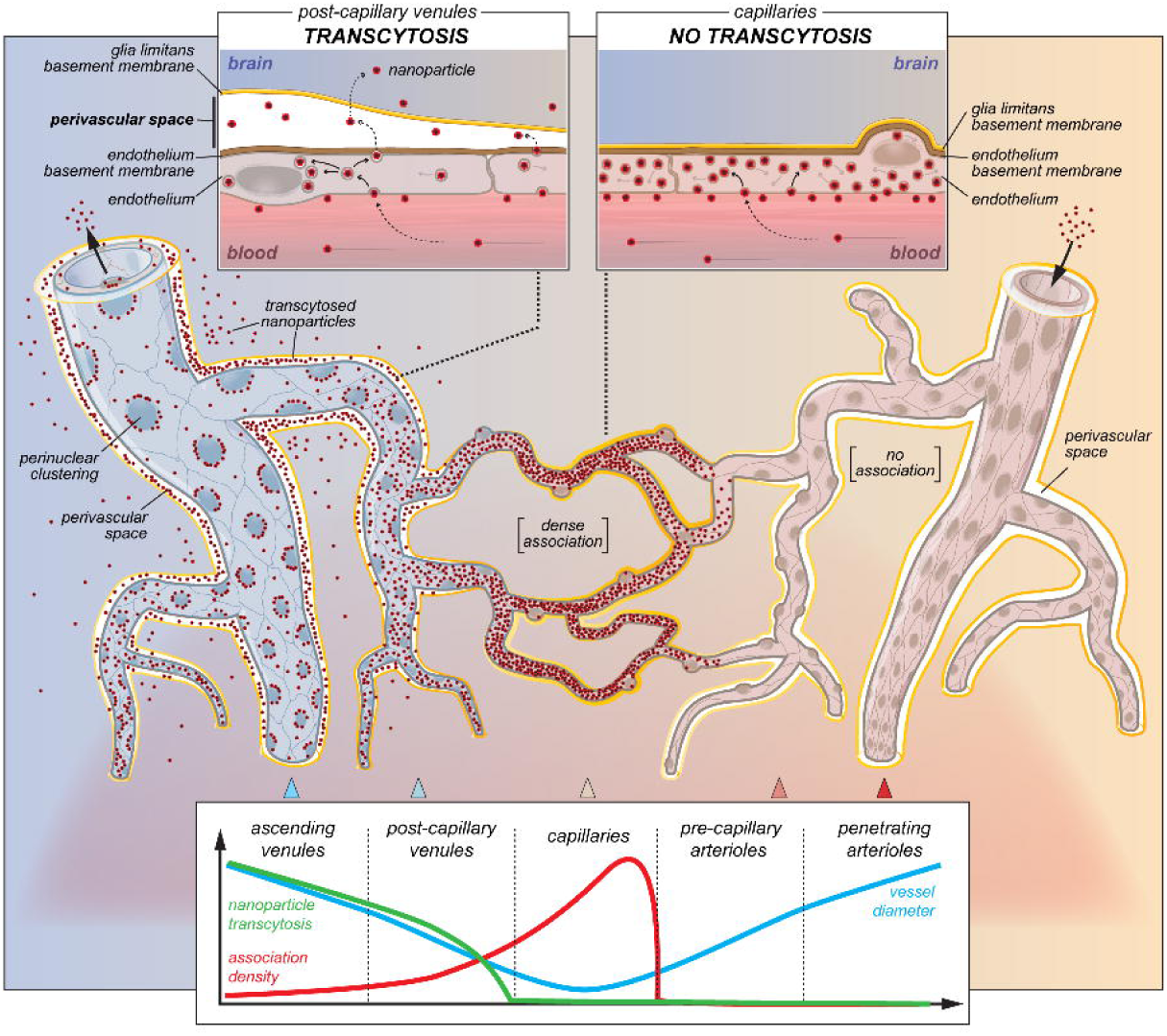
Results summary.

The association of nanoparticles to the endothelium rapidly declined from capillaries to venules, inversely proportional to the vessel diameter and consistent with the TfR expression profile (Ge et al., 2005; Jefferies et al., 1984). TfR is a recycling receptor with ∼10% of the total receptor pool present at the surface of BECs (van Gelder et al., 1995). This amount was sufficient to maintain steady and ongoing recruitment of TfR-targeted nanoparticles to the vessel wall without saturation of the receptor pool, even at the high densities in capillaries. Although anti-TfR antibodies readily cross the BBB (Johnsen et al., 2019b; Niewoehner et al., 2014; Yu et al., 2011), there is a fundamental disagreement as to whether significantly larger TfR-targeted nanoparticles actually enter the brain (Clark and Davis, 2015; Lindqvist et al., 2016; van Rooy et al., 2011; Wiley et al., 2013). Here, we provide direct evidence of TfR-mediated transcytosis of nanoparticles across the endothelial cells of the post-capillary venules, and challenge the concept that nanoparticles undergo transport across the BBB selectively at capillary endothelial cells.

The key question is why the highest density of associated nanoparticles in capillaries did not translate to the vascular segment of transcytosis. The lack of nanoparticle passage in capillaries can not be explained solely by impaired dissociation of antibody-functionalized nanoparticle from TfR during exocytosis (Couch et al., 2013; Yu et al., 2011), as transcytosis would also be absent at venules. Instead, we propose that the microanatomy of the tissue surrounding microvessels determines successful nanoparticle entry into the brain, specifically the size of the perivascular space and organization of the basement membrane. Large and post-capillary venules are surrounded by perivascular space located between the endothelium basement membrane and astrocyte glia limitans, but in contrast to venules, capillaries are devoid of perivascular space because both membranes are fused (Engelhardt et al., 2017; Thomsen et al., 2017). This principal anatomical brain feature is present in both murine and human brains (Zhang et al., 1990).

We propose that nanoparticles are more likely to transcytose via the route of the least resistance, i.e. into the perivascular space of venules; instead of entering the more restrictive compartment, i.e. the brain parenchyma at the capillary segment (Foley et al., 2012; Wardlaw et al., 2020). By analogy, translocation of blood-borne leukocytes across the BEC into the brain is also possible only at the microvascular segments surrounded by perivascular space capable of accommodating the cells, i.e. at post-capillary venules (Engelhardt et al., 2017). It also is unlikely that nanoparticles were transcytosed at capillaries and subsequently relocated downstream along the capillary wall into the perivascular space of venules due to the interstitial fluid flow. First, the hydraulic resistance around the capillaries is too high for interstitial fluid flow (Faghih and Sharp, 2018). Second, we detected no significant nanoparticle movement following laser extravasation to the brain tissue surrounding capillaries, compared to the rapid progression of nanoparticles extravasated from venules into the perivascular space (Movie S9). Third, if transcytosis occurred in capillaries, a significant fraction of nanoparticles of >100 nm diameter would likely become trapped in the protein meshwork of fused capillary basement membranes (Cabezon et al., 2015; Muldoon et al., 1999), but this we did not observe in chronic imaging experiments.

The major concern of the large nanoparticle-based drug delivery systems is the ability of transcytosed nanovehicles to progress within the extracellular space (ECS). *In vivo* diffusion experiments estimate the ECS to be in the range of 38–64 nm (Thorne and Nicholson, 2006), although recent super-resolution imaging revealed ECS clefts of ∼100 nm (Tonnesen et al., 2018). These dimensions may not restrict the movement of, e.g., antibodies, but may exclude significantly larger nanoparticles from entry and progression within the ECS (Nance et al., 2012; Thorne and Nicholson, 2006). Here, most nanoparticles were detected in perivascular areas, even after 2 days post-injection, which indicates high perivascular retention. However, we also observed nanoparticles at distances from post-capillary venules corresponding to the location of neuropil, indicating successful passage into the ECS. How ∼130-nm nanoparticles might travel within the ECS is unclear but appears to be highly dependent on the level of PEGylation (Nance et al., 2012). In the murine brain, neurons are located on average within short (∼15 *µ*m) distances from brain capillaries (Tsai et al., 2009). Although TfR may potentially act as an intraparenchymal target for neurons (Chen and Liu, 2012; Farias et al., 2015), we observed no cellular association patterns of the nanoparticles suggestive of binding to TfRs, which are enriched at the neuronal somata (Farias et al., 2015).

In contrast to microvessels embedded in the brain parenchyma, the vasculature in the dura lacks the BBB (Engelhardt et al., 2017; Weller et al., 2018). Contrary to a recent study (Lam et al., 2018), we detected no TfR-targeted nanoparticles in the dura. However, we did observe blood-borne immune cells that infiltrated the brain, carrying along nanoparticles that were hijacked from the bloodstream. Thus, nanoparticles could also enter the CNS in a manner independent of vesicular transcytosis. This phenomenon, in a similar manner to transcytosis, also occurred exclusively in venules. The leakage of large blood-borne macromolecules can reportedly occur via wedging of leukocytes between endothelial cells in venules but not in capillaries (Claudio et al., 1990). However, this phenomenon is associated with severe autoimmune reactions, and we did not detect the clustering of nanoparticles at BEC contact sites.

An important consideration is to what extent brain microanatomy might influence nanoparticle transcytosis compared to other BEC features along the vascular segments. First, colloidal glycocalyx at the BEC luminal side may potentially bury macromolecules within its matrix or repel negatively charged nanoconstructs, impairing endocytosis (Cheng et al., 2016; Gromnicova et al., 2016). Indeed, we detected that the internalization of nanoparticles was not an all-or-nothing phenomenon, and ∼30-40% total nanoparticles recruited from the blood circulation were restricted to the luminal side of BEC. Although enzymatic shedding of glycocalyx does not improve BEC uptake of positively charged nanoparticles (Gromnicova et al., 2016), we cannot exclude that glycocalyx might affect slightly negatively charged particles (Kutuzov et al., 2018), and in a vessel-type–dependent manner.

Second, our data show that after internalization, the nanoparticles exhibited similar movement dynamics regardless of vessel type and consistent with cytoskeleton-assisted transcytosis (Soldati and Schliwa, 2006). However, in contrast to capillaries, post-capillary and pial venules exhibited the preferential distribution of nanoparticles to perinuclear areas, which corresponds to the position of late endosomes in the transcytotic pathway that contain components directed for degradation (De Bock et al., 2016; Grant and Donaldson, 2009). Although this suggests a more restrictive intracellular barrier for the trafficking of nanoparticles in venules than in capillaries, it supports the notion that despite segmental differences in RMT, it is ultimately the microanatomy of the brain surrounding microvessels that plays a decisive role for successful transcytosis.

Lastly, the vessel-dependent manner in which the brain handles TfR-targeted nanoparticles may potentially explain conflicting reports on the delivery of other TfR-targeted entities, such as antibodies, where the robust ability to penetrate the BBB is often opposed by evidence for the same type of entity indefinitely trapped within the capillaries (Friden et al., 1991; Moos and Morgan, 2001). Our findings may be of relevance for other than TfR-targeting approaches and potentially explain why other targeting ligands e.g. COG133, angiopep-2, or CRM197 that performed successfully in capillaries *in vitro*, failed to enhance nanoliposome uptake *in vivo* (van Rooy et al., 2011). Our methodological platform is first to describe the nanoparticle fate in the living brain at nanoscale resolution in live, and also, awake animals. It is suitable for examining the potential effects of disease states on large molecule therapeutics delivery, e.g. during stroke-induced imbalance in transcytosis (Knowland et al., 2014), reduction of perivascular spaces (Mestre et al., 2018), or ECS changes in edema or during brain activity (Tonnesen et al., 2018). Our findings may also help to avoid pitfalls in design of drug delivery systems, e.g. in Alzheimer disease, where Aβ is deposited along pericapillary and periarteriolar spaces (Tarasoff-Conway et al., 2015), rather than in venular perivascular space, which is the locus of nanoparticle transcytosis.

In summary, we reveal the transcytosis pathway across the BBB of targeted clinically relevant nanoparticles. Our identification of post-capillary venules as the key site for transcytosis may aid efforts to develop efficient therapeutic approaches for drug delivery to the CNS. As an accessible part of the vascular tree, post-capillary venules may be poised to serve as a preferred route for macromolecule and nanoparticle delivery across the BBB.

## ACKNOWLEDGMENTS

This study was supported by Lundbeck Foundation Research Initiative on Brain Barriers and Drug Delivery (#R392-2018-2266); Det Frie Forskningsråd (#0602-01965B); Nordea Foundation Grant to the Center for Healthy Aging (#114995). We thank Nikolay Kutuzov for feedback on data analysis, Serhii Kostrikov for his assistance in cisplatin experiments, and Kjeld Møllgård for his comments on the manuscript.

## AUTHOR CONTRIBUTIONS (CRediT taxonomy)

Conceptualization, Kr. Ku., Ka. Kr. and M. J. L.; Methodology, Kr. Ku. and Ka. Kr.; Formal Analysis, Kr. Ku.; Investigation, Kr. Ku., Ka. Kr., K.B.J., M. A. L., and M. L.; Resources - T.M., T. L. A. and M. J. L.; Writing – Original Draft, Kr. Ku.; and M. J. L.; Writing – Review & Editing, Kr. Ku., Ka. Kr., K.B.J., T.M., and M.J.L.; Visualization, Kr. Ku.; Funding Acquisition, T.M., T. L. A. and M. J. L.; Supervision M. J. L.

## Methods

### Animals

All animal experiments were approved by the Danish National Ethics Committee and followed ARRIVE guidelines. We used wild-type C57Bl/6 mice, age 5–7 months (23–31 g) and age-matched (25–32 g) homozygous Tg(TIE2GFP)287Sato/J transgenic reporter mice (Tie2-GFP mice, #003658, The Jackson Laboratory) expressing GFP under the endothelial-specific receptor tyrosine kinase (Tie2) promoter (Motoike et al., 2000).

### Animal preparation for acute imaging

Surgery was performed as previously described, with minor modifications (Kucharz and Lauritzen, 2018). Briefly, animals were anesthetized by intraperitoneal (i.p.) injection of xylazine (10 µg/g_animal_) and ketamine (60 µg/g, then 30 µg/g_animal_, at 20– 25 min intervals). A tracheotomy was performed for mechanical respiration (180–220 μL volume at 190–240 strokes/min; MiniVent Type 845, Harvard Apparatus) with O_2_-supplemented air (1.5–2 mL/min). Two catheters were inserted, one into the left femoral artery for injection of compounds and nanoparticles, and for monitoring mean arterial blood pressure (MABP; Pressure Monitor BP-1, World Precision Instruments), and the other into the femoral vein for anesthesia infusion during imaging. The animal was turned to the prone position, the scalp was removed, the periosteum was removed with a FeCl_3_-soaked cotton bud, and the exposed skull was glued (Loctite Adhesives) to a custom-made metal head plate. A craniotomy was performed over the right somatosensory cortex (3 mm lateral, 0.5 mm posterior to bregma; Ø=4 mm; 4500 rpm dental drill). The bone flap was carefully lifted, the dura removed, and 1% agarose (type III-A, Sigma-Aldrich) in artificial cerebrospinal fluid (aCSF; in mM: NaCl 120, KCl 2.8, Na_2_HPO_4_ 1, MgCl_2_ 0.876, NaHCO_3_ 22, CaCl_2_ 1.45, glucose 2.55, pH=7.4) was applied on the brain surface. An imaging coverslip (∼4×4 mm, 0.08-mm thick; Menzel-Gläser) was positioned onto the craniotomy, leaving a ∼0.5-mm gap for glass microelectrode insertion. The animal was transferred to the imaging stage, and the anesthesia was changed to continuous infusion of α-chloralose (50 mg/kg BW per hour) via an intravenous catheter.

Mice were allowed to rest for 25 min before the imaging procedures. Prior to imaging, a ∼50 µL blood sample was collected via the arterial catheter for blood gas evaluation (ABL, Radiometer), and the respiration rate and volume were adjusted if necessary. To ensure physiological conditions, we monitored end-tidal CO_2_ levels and MABP, and body temperature was maintained at 37°C using a rectal thermistor-regulated heating pad.

### Animal preparation for awake/chronic awake imaging

The surgery was performed as previously described, with minor modifications (Holtmaat et al., 2009). Briefly, 4 h prior to the surgery, the animals were injected with dexamethasone (4.8 mg/g BW; Dexavit, Vital Pharma Nordic). The anesthesia was induced with 5% isoflurane (ScanVet) in O_2_-supplemented air (10%). Eyes were lubricated with eye ointment (Viscotears, Novartis), and the animal’s head was shaved and mounted onto a stereotactic frame. The body temperature was maintained during all steps of the surgery at 37°C using a rectal thermistor-regulated heating pad. Surgery was performed in an aseptic environment with heat-sterilized surgical tools. The shaved skin was disinfected with chlorhexidine/alcohol (0.5%/74%; Kruuse). Next, carprofen (5 mg/kg BW; Norodyl, Norbrook), buprenorphine (0.05 mg/kg BW; Temgesic, Indivior), and lidocaine (100 µL 0.5%) were subcutaneously injected under the scalp. The anesthesia was reduced to 1.8%–2.0% isoflurane, the scalp was removed, and the bone surface was cleaned from the periosteum with an ultrasonic drill (Piezosurgery, Mectron). A craniotomy was performed over the right somatosensory cortex (2 mm lateral, 0.5 mm posterior to bregma; Ø=3 mm), the bone flap was carefully lifted, and the exposed brain temporarily covered with a hemostatic absorbable gelatin sponge (Spongostan®, Ferrosan, Denmark) pre-wetted with ice-cold aCSF. The cranial opening was filled with aCSF, then sealed with an autoclave-sterilized round imaging coverslip (Ø=4 mm, 0.17-mm thick; Laser Micromachining LTD). The rim of the coverslip was secured with a thin layer of Vetbond (3M), and a lightweight stainless steel head plate (Neurotar) was positioned on the top of the skull in alignment with the cranial window. The skull was coated with adhesive resin cement (RelyX Ultimate, 3M) to secure the exposed bone, including skin incision rim, and to firmly attach the metal plate to the head. Next, the animals were transferred onto a pre-warmed heating pad to wake from anesthesia (∼5 min) in a cage supplemented with pre-wetted food pellets for easy chow and hydration.

Post-operation care consisted of subcutaneous injections of Temgesic (3 h) and Rimadyl (24 and 48 h post-surgery, doses as before). Animal welfare was closely monitored during the 7 days of post-surgical recovery and subsequent imaging training procedures. All animals underwent recurrent 30-min/day training sessions before the imaging to gradually habituate to the mobile cage system (Neurotar) with sugar-supplemented water as a reward (∼14 days training). Given that no catheters were mounted in chronically imaged animals, the nanoparticles were injected retroorbitally 10 days post-surgery during brief (∼2 min) isoflurane anesthesia (5%). This administration route was preferential to, e.g., tail vein injections, because it provided better control over the injectant volume. The imaging sessions never exceeded 45 min.

### Nanoparticle preparation

1,2-Distearoyl-*sn*-glycero-3-phosphocholine (DSPC), ovine cholesterol, 1,2-distearoyl-*sn*-glycero-3-phosphoethanolamine-N-[methoxy(polyethylene glycol)-2000] ammonium salt (DSPE-PEG2k), and 1,2-distearoyl-*sn*-glycero-3-phosphoethanolamine-N-[maleimide(polyethylene glycol)-2000] ammonium salt (DSPE-PEG2k-maleimide) were purchased from Avanti Polar Lipids (Alabaster, AL, USA). The stealth nanoparticles were prepared to consist of DSPC/cholesterol/DSPE-PEG2k (molar composition: 56.3:38.2:5.5), and the antibody-functionalized nanoparticles to consist of DSPC/cholesterol/DSPE-PEG2k/DSPE-PEG2k-maleimide (molar composition: 56.3:38.2:5:0.5). The fluorescent nanoparticles were supplemented with 0.5 mol% of 1,2-dipalmitoyl-*sn*-glycero-3-phosphoethanolamine labeled with Atto488 (Atto488 DPPE) or Atto550 (Atto550 DPPE; Atto-Tec, Siegen, Germany). To obtain lipid powder mixtures of the above compositions, the constituent lipids were dissolved in tert-butanol (Sigma-Aldrich, St. Louis, MO, USA)/Milli-Q water solution (9:1) and heated to 40–50°C to ensure complete dissolution. The lipid solutions were then plunge-frozen in liquid N_2_ and lyophilized overnight to remove the solvent (ScanVac CoolSafe lyophilizer, LaboGene, Allerød, Denmark).

### Nanoparticle fluorescent tagging

To obtain fluorescently labeled nanoparticles, the lyophilized lipids were hydrated in 70°C phosphate-buffered saline (PBS; 10 mM phosphate, 137 mM sodium chloride, 2.7 mM KCl, pH 7.4; tablets from Sigma-Aldrich) to a 50 mM lipid concentration. The lipid suspensions were vortexed seven times in 5-min intervals, then subjected to five freeze-thaw cycles by alternate placement in a liquid N_2_ and a 70°C water bath. Next, the lipid suspensions were extruded 21 times through a 100-nm polycarbonate filter (Whatman, GE Healthcare) at 70°C using a mini-extruder (Avanti Polar Lipids) to form nanoparticles.

### Nanoparticle targeting

We used a high-affinity (K_D_=6 nM) monoclonal anti-TfR antibody clone RI7217 to functionalize nanoparticles (Johnsen et al., 2018). The antibody was produced in-house using the hybridoma technique at Laboratory for Neurobiology, Aalborg University, Denmark. The antibody specificity was previously determined using surface plasmon resonance (Johnsen et al., 2018). To functionalize the nanoparticles with either RI7217 or a rat isotype IgG control (Thermo Fisher Scientific, Waltham, MA, USA), we prepared solutions of 8 mg/mL antibody in borate buffer (100 mM borate, 2 mM EDTA, pH 8.0; all Sigma-Aldrich). The antibody concentrations were determined from the absorbance at 280 nm (NanoDrop 2000c spectrophotometer, NanoDrop Products, Thermo Fisher Scientific) using mass extinction coefficients of 1.3 (mg/mL)^-1^ cm^-1^ and 1.5 (mg/mL)^-1^ cm^-1^ for RI7217 and IgG isotype, respectively, determined in a separate micro-BCA experiment. Traut’s reagent (Thermo Fisher Scientific) was added to a reagent-to-antibody molar ratio 10:1 in Protein LoBind tubes (Eppendorf, Hamburg, Germany), and the solutions were incubated for 1 h at room temperature under constant shaking at 500 rpm. Using Amicon Ultra-4 30 kDa centrifugal filter units (Merck, Darmstadt, Germany), we transferred the thiolated antibodies to PBS and determined their concentration using the NanoDrop 2000c as described above. Next, we added 1.05 mg of newly prepared thiolated antibodies to 700 µL of newly prepared nanoparticles (lipid concentration ∼35–40 mM) in Protein LoBind tubes and replaced the air phase in the tubes with N_2_. The samples were then incubated for 24 h at room temperature under constant shaking at 500 rpm, allowing the thiolated antibodies to couple to the maleimide groups the surface of the nanoparticles. The antibody-functionalized nanoparticles were separated from unbound antibodies using a Sepharose CL-4B (GE Healthcare, Little Chalfont, UK) size-exclusion chromatography column eluted with PBS (dimensions, 1.5×20 cm; flow rate, 1 mL/min). The recovered nanoparticles were pooled in Amicon Ultra-4 100 kDa centrifugal filter units (Merck) and concentrated by centrifuging at 2000 ×*g* until the lipid concentration was increased to 30–40 mM.

### Nanoparticle cisplatin loading, targeting and detection

To prepare cisplatin-loaded nanoparticles, *cis*-diammineplatinum(II) dichloride (cisplatin; Sigma-Aldrich) was dissolved in PBS to a nominal concentration of 8.5 mg/mL (Peleg-Shulman et al., 2001). The solutions were magnetically stirred for 1 h at 70°C and subsequently left at room temperature for 15 min, allowing undissolved cisplatin crystals to precipitate. The supernatants were transferred to new vials and magnetically stirred while being heated to 70°C. The solutions were then added to lyophilized lipids, resulting in 50 mM lipid suspensions that were magnetically stirred for 1 h at 70°C and extruded as described above for the fluorescently labeled nanoparticles. The samples were cooled to room temperature to allow any residual cisplatin crystals to precipitate, and the supernatants were run on a Sepharose CL-4B size-exclusion chromatography column eluted with PBS (dimensions 1.5×20 cm, flow rate 1 mL/min) to remove free cisplatin. The recovered nanoparticles were concentrated using Amicon Ultra-4 100 kDa centrifugal filter units by centrifuging at 2000 ×*g*.

To prepare antibody-functionalized cisplatin-loaded nanoparticles, the antibodies were thiolated as described above. Then, 0.5 mg of newly prepared thiolated antibody was added to 700 µL of newly prepared cisplatin-loaded nanoparticles (lipid concentration ∼13 mM) in a Protein LoBind tube. The nanoparticles were finally incubated, recovered, and concentrated as described above for the fluorescently labeled antibody-functionalized nanoparticles.

Brain uptake of cisplatin was measured using ICP-MS as recently described (Johnsen et al., 2019a).

### Nanoparticle properties *in vitro*

The phosphorus concentrations of the liposome stock solutions were determined using inductively coupled plasma mass spectrometry (iCAP Q ICP-MS, Thermo Fisher Scientific). We estimated phospholipid concentrations by subtracting the phosphorus background of the PBS buffer and estimated total lipid concentrations by dividing the phospholipid concentrations by 0.618, taking into account that cholesterol does not contain phosphorus. For the cisplatin-loaded nanoparticles, we also used ICP-MS to determine the platinum concentrations. The size of the nanoparticles (dissolved in PBS) was measured using dynamic light scattering, and the zeta potential of the nanoparticles in phosphate-glucose buffer (10 mM phosphate, 280 mM glucose, pH 7.4; reagents from Sigma-Aldrich) was measured using mixed measurement mode phase analysis light scattering (Zetasizer Nano ZS, Malvern Instruments, Malvern, UK). The antibody conjugation level on the functionalized nanoparticle was determined using the micro-BCA assay (reagents purchased from Thermo Fisher Scientific), performed by incubating samples (including bovine serum albumin [BSA] standard samples) for 1 h in a 60°C water bath and then transferring them to a 96-well plate to measure their absorbance at 562 nm using a Spark multimode microplate reader (Tecan, Männedorf, Switzerland). To account for the small contribution of lipids in the micro-BCA assay (Kessler and Fanestil, 1986; Kristensen et al., 2019), we also performed the micro-BCA on non-functionalized nanoparticles, which allowed for the subtraction of the lipid contribution to determine the amount of antibody conjugated to the nanoparticles. The hydrodynamic diameter (D_h_) of RI7-functionalized Atto 550-tagged (RI7-L-A550) and Atto 488-tagged (RI7-L-A488) nanoparticles was in the range of d_h_=∼135–140 nm (Table S1) and comparable to other TfR-targeted clinically relevant formulations (Johnsen and Moos, 2016; Lam et al., 2018). Both RI7-L-A550 and RI7-L-A488 had a low polydispersity index (≤0.13), indicating high size homogeneity. Assuming the nanoparticles contained on average ∼2.5×10^5^ lipids, the conjugation level of 30 g/mol_lipid_ corresponded to ∼50 antibodies per nanoparticle (Kristensen et al., 2019).

### Nanoparticle systemic interactions *in vivo*

In contrast to other TfR ligands, RI7217 does not compete with endogenous transferrin, and TfR vascular expression is highly specific to the brain (van Rooy et al., 2011). We observed no pathological changes in exhaled CO_2_ levels, mean arterial blood pressure MABP, or brain activity after nanoparticle administration in acute experiments (Figure S1). Furthermore, we detected no atypical behavior, weight loss, or signs of neuroinflammation in animals monitored up to 48 h post-injection in chronic imaging experiments.

### Other fluorescent probes

FITC-dextran (MW 10 kDa, 0.5%, Sigma-Aldrich), TRITC-dextran (MW 65 kDa, 1%, Sigma-Aldrich), or bovine serum albumin Alexa Fluor 488-conjugate (BSA-Alexa 488, 1%, Invitrogen) was administered as a single bolus injection (50 µL) via a femoral arterial catheter. All were dissolved in sterile saline and administered subsequently to nanoparticles. In addition to delineating a vessel lumen, lack of extravascular leak of dyes indicated preserved BBB structural integrity after the microsurgery.

### Imaging setup

*In vivo* two-photon imaging was performed with a SP5 upright laser scanning microscope (Leica Microsystems) coupled to MaiTai Ti:Sapphire laser (Spectra-Physics). The images were collected using a 20× 1.0 water-immersion objective. The fluorescence signal was split by FITC/TRITC filter and collected by two separate multi-alkali photomultipliers after 525–560 nm and 560–625 nm bandpass filter (Leica Microsystems). The fluorophores were excited at 870 nm with the 14 mWatt output power at the sample. The images were collected using LAS AF v. 4.4 (Leica Microsystems) in 16-bit color depth and exported to ImageJ for further analysis (v. 1.52a; NIH). 3D reconstructions were performed via volume rendering in Amira v. 6 (FEI Visualization Sciences Group).

### Surface density calculation

To assess the spatio-temporal properties of nanoparticles association to the endothelium, we monitored the association of nanoparticles for 2 h after injection with respect to all cerebral vessel types using hyperstack (4-dimensional) imaging. Data were recorded as a series of Z-stacks in bidirectional scanning mode with triple frame averaging, from 387.5 μm×387.5 μm area (2048×2048 pixel resolution) and 144 μm depth span (Z-step size 2.50 μm) with 7.5-min intervals between consecutive Z-stacks. The cerebral microvessels were classified as pial arterioles or venules, penetrating arterioles, post-capillary venules, ascending venules, or capillaries based on their morphology and second harmonics generation (Grubb et al., 2020; Janiurek et al., 2019). The nanoparticles were counted from all vessels in the field of view, with each individual vessel followed over time. The vessel surface area was calculated from vessel diameter delineated by FITC-dx or TRITC-dx signal and the length of the vessel measured in 3D. The association density was obtained from a nanoparticle count per corresponding vessel surface area [nanoparticles/µm^2^].

### Subcellular distribution mapping

We imaged the surface of the vessels, i.e., a ∼5 µm planar optical section aligned with the vessel circumference. When measuring distances from the nucleus geometric center to nanoparticles, we set the distance cut-off point to 11 µm to exclude the nanoparticles that belonged to neighboring endothelial cells and to avoid distribution bias due to the non-concentric spindle-like geometry of endothelial cells. In addition, we excluded nanoparticles located in line from the geometric center towards the vessel wall, where the cut-off distance exceeded the vessel boundary. We took this step to minimize the effect of the vessel curvature on the estimation of the distance.

### MSD analysis

To characterize nanoparticle movement dynamics, we used the mean square displacement (MSD) analysis (Weimann et al., 2013). Time-lapse recordings were collected in bidirectional scanning mode from 387.5 μm×387.5 μm area (2048×2048 pixel resolution) for 30 min at 30-s intervals (60 data timepoints). Each nanoparticle trajectory was manually traced, treating the fluorescence radial symmetry center as a nanoparticle location coordinate. The planar (*x,y*) trajectories were projected to a vector *(v)* aligned with a vessel symmetry axis and with the direction of the blood flow. For every trajectory, the displacements in *v* coordinate in time *t* were extracted for each multiplier of the smallest resolved time interval *d* (i.e., *t=d, t=*2*d, t=*3*d*…*t=i*d*), where *i=*60 timepoints and *d*=30 s. In simple model systems, the significant deviation of MSD(t) from linearity with the increase of *t* indicates a non-stochastic (directional) movement component (Weimann et al., 2013). We assessed MSDv(*t*) linearity with the least squares linear regression fit weighted by the inverse of data point variance (Michalet, 2010), followed by Wald–Wolfowitz runs test.

### Electrophysiological recordings

Electrocortical brain activity (ECoG) was recorded via a heat-pulled borosilicate glass electrode containing an Ag/AgCl filament and filled with aCSF (electrode tip Ø, 2–3 μm; inner Ø, 0.86 mm; outer Ø, 1.5 mm; Sutter Instrument; resistance 1.5–2.0 MΩ). The electrode was inserted under the glass coverslip ∼50 μm into the cerebral cortex, and the reference electrode was positioned in the neck muscle. The total electrical signal was filtered (3000 Hz low-pass filter), then amplified 10× (AP311 analog amplifier; Warner Instruments), and the alternate current-ECoG component (i.e., spontaneous brain activity) was obtained after further 100× amplification and 0.5 Hz high-pass filter (NL 106 analog amplifier and NL 125/126 analog filter, NeuroLog). Analog data were digitized (Power 1401, CED) at 20 kHz. For the exhaled CO_2_, MABP (the raw readout) was collected. All data were recorded in Spike2 software (v. 7.02a; CED).

### Statistical analysis

Following Pearson’s normality test, either an unpaired two-tailed Student’s t-test with Welch’s correction (normally distributed data) or Mann–Whitney test (non-normally distributed data) was used. All statistical analyses were performed in Prism v.8.2 (GraphPad). The sample size was selected based on our previous two-photon *in vivo* experiments (Kucharz and Lauritzen, 2018). Data was plotted in Prism v.8.2. or in OriginPro 2018 (OriginLab Corporation).

### Exclusion criteria

Prior to injection of nanoparticles, all animals with abnormal blood pressure (<50 mmHg), abnormal brain ECoG activity, or significant (>2 µm/min) brain movement in *x, y*, or *z* coordinates were excluded from analysis (3 of 48 animals).

## SUPPLEMENTAL FIGURES LEGENDS

**Figure S1. No adverse systemic effects on physiology**.

**A)** Blood-circulating leukocytes with sequestered nanoparticles (arrowheads). See also Movie S2.

**B)** Leukocytes with sequestered nanoparticles preserve their rolling and endothelium adherence properties.

**C)** The uptake of nanoparticles is unlikely driven by the RI7217 moiety, as it is also present e.g. for fluorescently labeled albumin (BSA-Alexa488).

**D)** No significant effect of nanoparticles on brain activity (electrocorticogram, *ECoG*), exhaled CO_2_ levels (exCO_2_), and mean arterial blood pressure (*MABP*), all measured simultaneously with imaging.

**Figure S2. No nanoparticle association without TfR-targeting moiety**.

**A-B)** No association of stealth Atto 550-tagged nanoparticles (Sth-L-A550) to pial vessels and capillaries.

**C-D)** No association of isotype IgG Atto 550-tagged nanoparticles (IgG-L-A550) to pial vessels and capillaries.

**Figure S3. RI7-L-A488 follows RI7-L-A550 spatio-temporal distribution**.

**A–D)** High resolution *in vivo* images of RI7-L-A488 nanoparticles (green) 2 h post-injection. Co-injected circulating TRITC-dx delineates vessel lumen (red). Liposomes are readily present at pial, ascending venules, post-capillary venules, and capillaries *in vivo*.

**E)** No association of isotype IgG A488-tagged nanoparticles with pial vessels and capillaries.

**F-G)** Association of RI7-L-A488 over time *in vivo*. Nanoparticles associated most rapidly to endothelium of capillaries and most slowly to large venules. Surface density=# of nanoparticles per µm^2^ vessel wall area. Inset illustrates vessel type classification.

*mV*=main pial venule; *pV=*pial venule; *ascV=*ascending venule; *pcV=*post-capillary venule; *cap=*capillaries; *pA=*pial arteriole.

**Figure S4. Two-photon imaging of brain microvasculature endothelium *in vivo***.

**A)** Projected Z-stacks from Tie2-GFP mouse cortex showing brain microvasculature endothelium *in vivo*. The high-resolution scan details the subcellular morphology of arterioles and venules.

*pV=*pial venule; *ascV=*ascending venule; *pcV=*post-capillary venule; *cap=*capillaries; *penA=* penetrating arteriole; *pA=*pial arteriole; *nuc*=nucleus; cs=cell boundaries/endothelium contact sites.

## SUPPLEMENTAL MOVIES LEGENDS

**Movie S1**. Blood-circulating nanoparticles show high fluorescence stability over time. Time is relative to the time of nanoparticle injection.

**Movie S2**. A small fraction of nanoparticles is sequestered by circulating leukocytes.

**Movie S3**. Nanoparticle association to the brain microvasculature over time. Concurrent imaging of RI7-L-A550 nanoparticles (red) with FITC-dextran (FITC-dx, green). Time is relative to the time of nanoparticle injection.

**Movie S4**. Time-lapse recording of nanoparticle (RI7-L-A550) movement in the brain endothelial cells. The endothelium is visible in green (Tie2-GFP). *Left panels*: pial venule; *right panel*: post-capillary venule.

**Movie S5**. Time-lapse recording of nanoparticle (RI7-L-A550) motility at the capillary segment with examples of nanoparticle tracing. Circles denote individual nanoparticles.

**Movie S6**. Nanoparticles exhibit movement, even at stalled capillaries.

**Movie S7**. Right panel: nanoparticle (RI7-L-A550, red) transcytosis into the perivascular space at the level of post-capillary venules. Left panel: no nanoparticle transcytosis in capillaries. The vessel lumen is delineated by circulating FITC-dextran (FITC-dx, green).

**Movie S8**. Leukocyte entry into the brain with previously sequestered nanoparticles (RI7-L-A550, red) in the blood circulation. Vessel lumen is delineated by circulating FITC-dextran (FITC-dx, green).

**Movie S9**. No significant nanoparticle movement following laser-extravasation from the blood to the brain at capillaries. In contrast, rapid progression in the brain of nanoparticles extravasated from venules. Lines denote vessel lumen boundaries.

