## Supplementary Table 1 for "Post-capillary venules is the locus for transcytosis of therapeutic nanoparticles to the brain"

### SUPPLEMENTAL TABLES:

| | Nanoparticle | $D_h$ [nm] | PDI | Zeta potential [mV] | Fluorescence [a.u.] | Antibody [g/mol lipid] |
|---|---|---|---|---|---|---|
| Fluorescent nanoparticles | Sth-L-A488 | 120 | 0.04 | -9 | 17000 | 0 |
| | Sth-L-A550 | 116±5 | 0.02±0.02 | -7±1 | 14000±7000 | 0 |
| | RI7-L-A488 | 139 | 0.13 | -10 | 16000 | 29 |
| | RI7-L-A550 | 137±4 | 0.12±0.01 | -7±1 | 10000±3000 | 35±4 |
| | IgG-L-A488 | 135 | 0.10 | -9 | 14000 | 23 |
| | IgG-L-A550 | 136±5 | 0.10±0.03 | -7±1 | 9000±2000 | 31±20 |
| | Nanoparticle | $D_h$ [nm] | PDI | Cisplatin [g/mol lipid] | | Antibody [g/mol lipid] |
| Cisplatin-loaded nanoparticles | Sth-L-Pt | 118 | 0.03 | 15 | | 0 |
| | RI7-L-Pt | 161 | 0.16 | 13 | | 66 |
| | IgG-L-Pt | 168 | 0.13 | 13 | | 89 |

**Table S1. Related to Figure 1. Characteristics of nanoparticles.** $D_h$=hydrodynamic diameter; PDI=polydispersity index; a.u.=arbitrary unit. Data are averages±SD.
