## Supplementary figures and images for "Post-capillary venules is the locus for transcytosis of therapeutic nanoparticles to the brain"

### Supplementary Figure 1

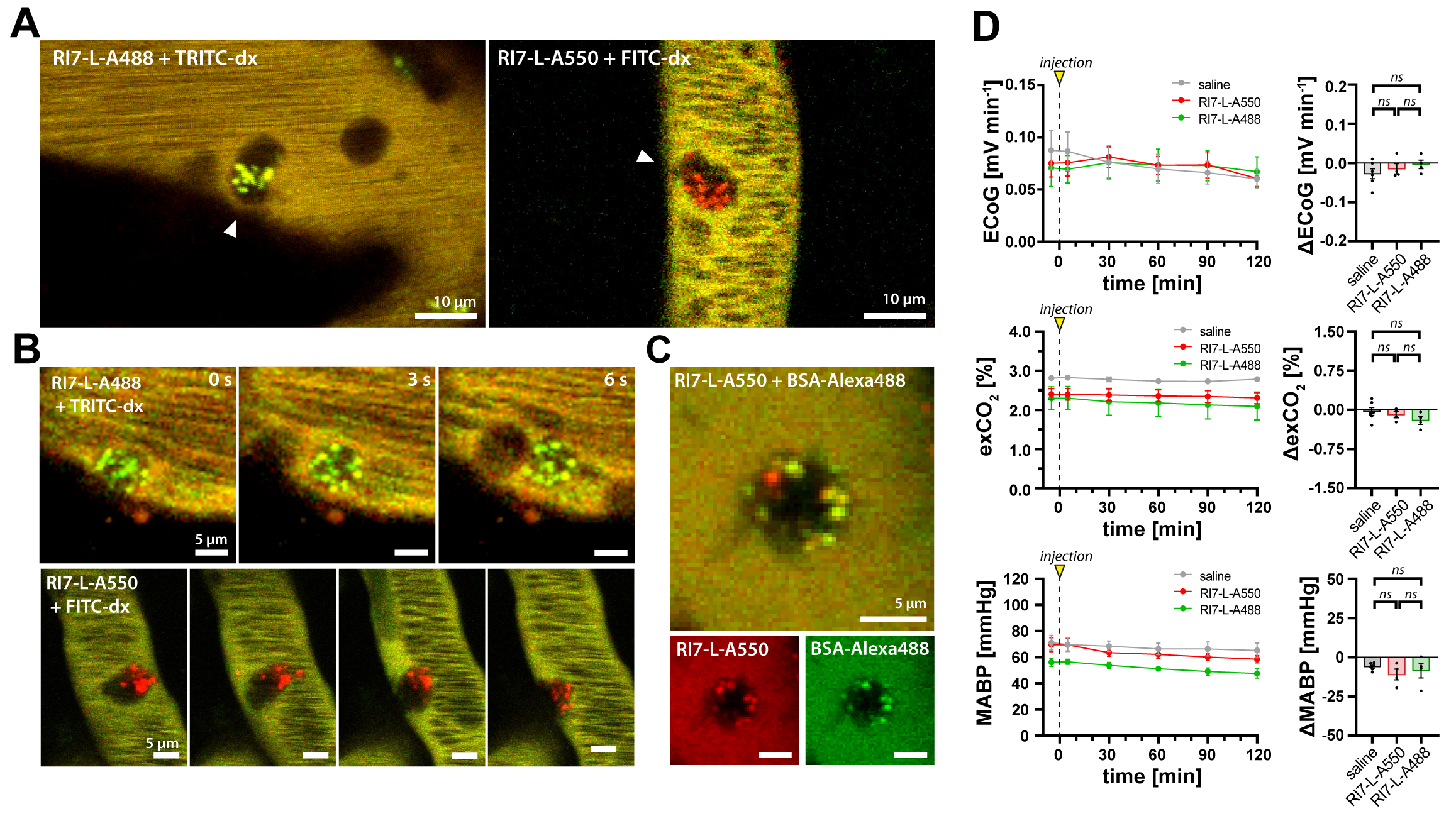

### Supplementary Figure 2

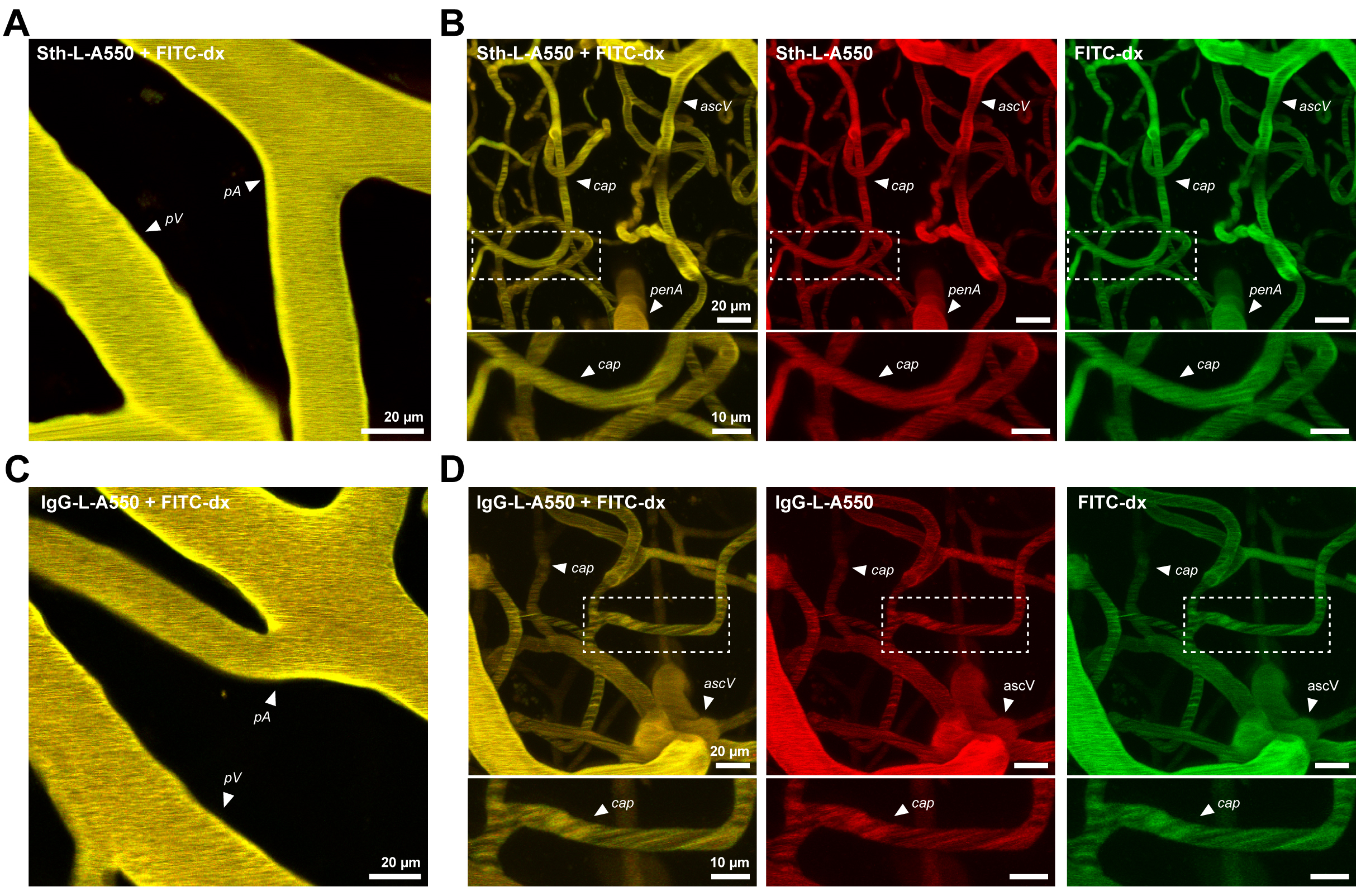

### Supplementary Figure 3

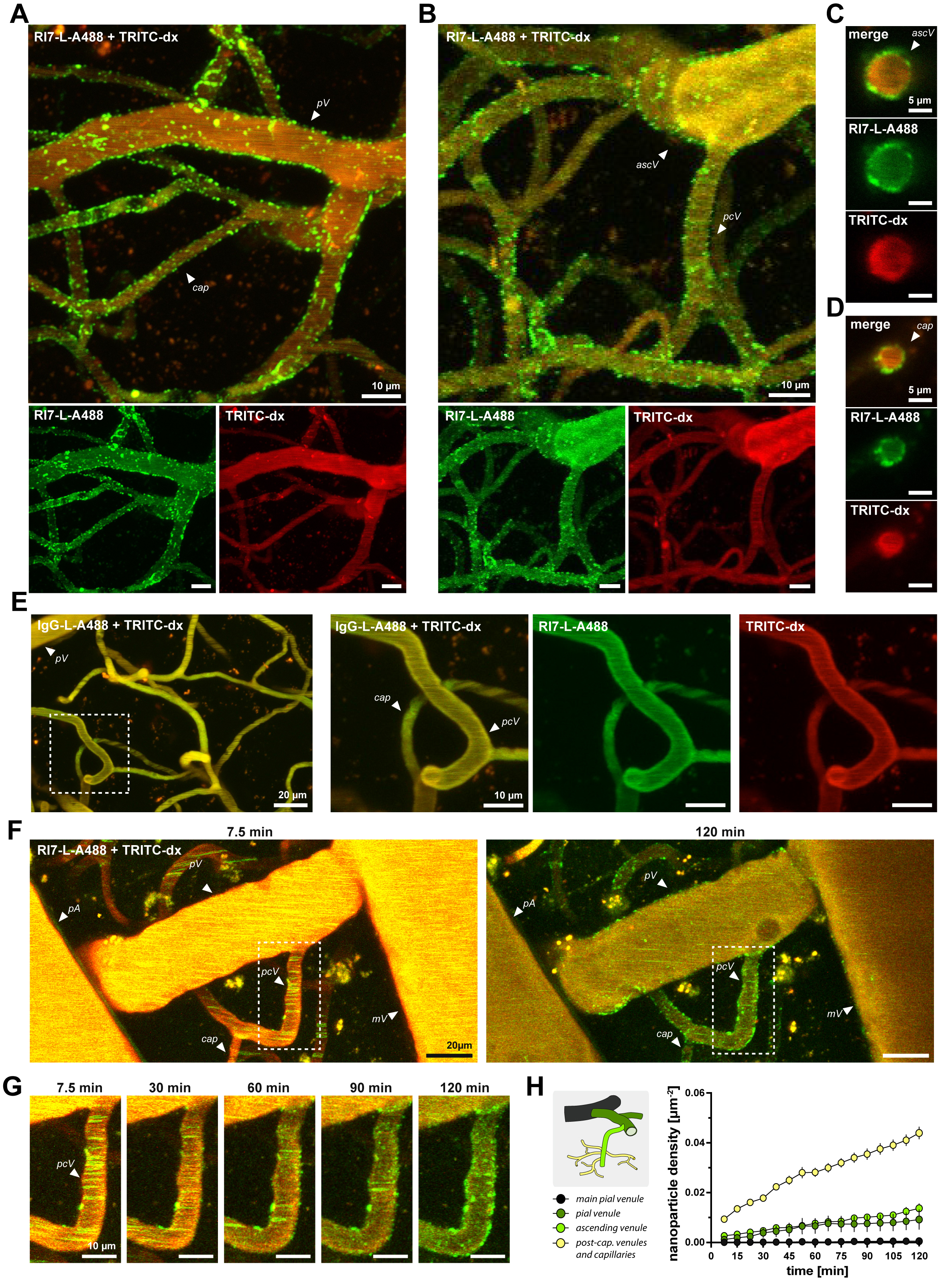

### Supplementary Figure 4

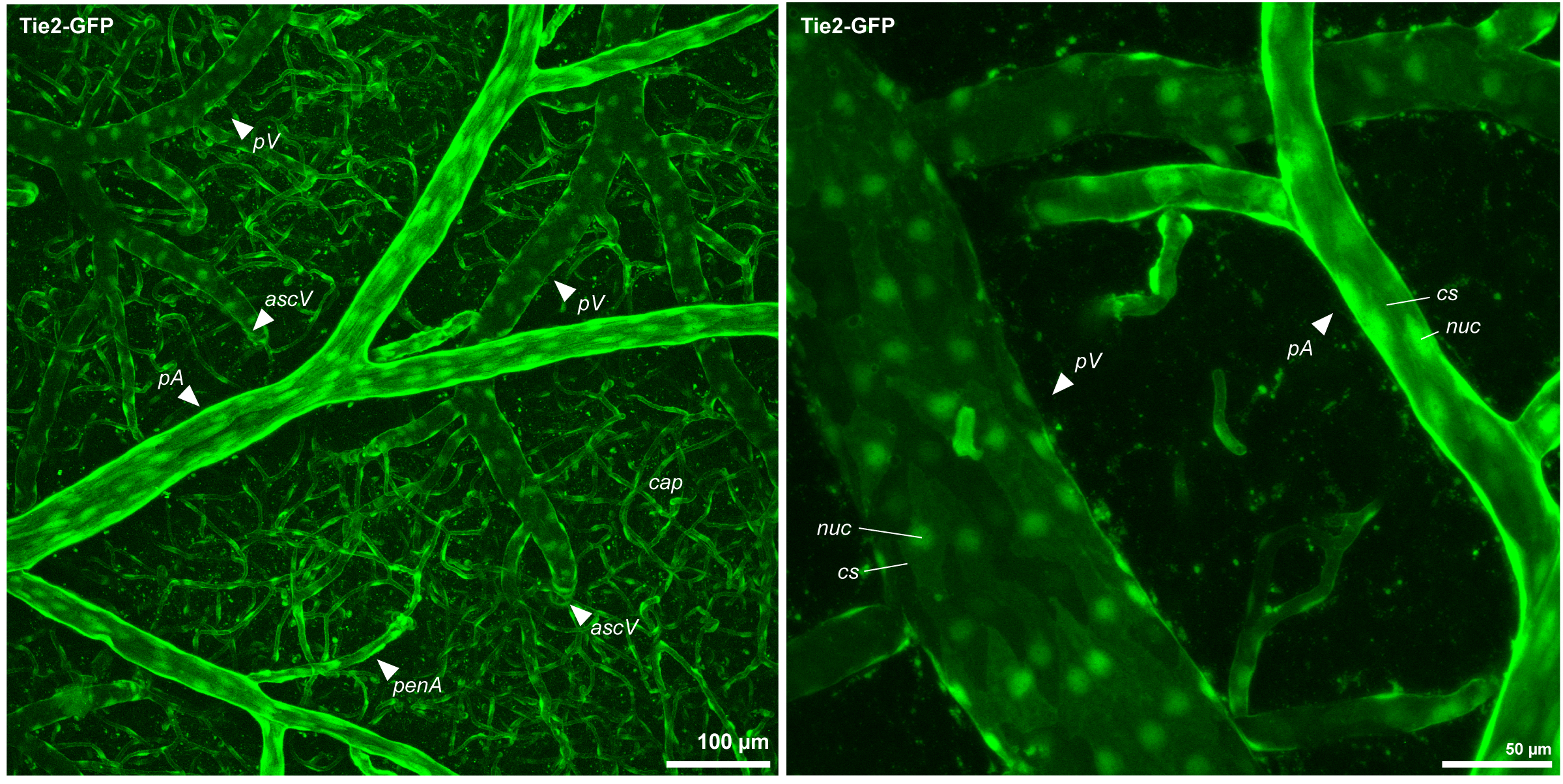
